# Imitation performance in primary school children

**DOI:** 10.1101/2022.04.27.489190

**Authors:** Giovanni Ottoboni, Alessio Toraldo, Riccardo Proietti, Angelo Cangelosi, Alessia Tessari

## Abstract

We studied the development of imitation ability in a cohort of 6-11-year-old children (N=174) with specific attention to error types and their cognitive interpretation. Participants imitated meaningless actions as if they were in front of a mirror (specularly). Actions varied across three levels of complexity (movements of a single limb, arm and leg of the same body side, or arm and leg of opposite sides). Overall performance improved with age. Among the most frequent error categories, ‘side’ errors (movement imitated with the left instead of the right limb or vice-versa) paradoxically increased with age (from 9 years). Still, their sensitivity to complexity decreased with age. Thus, encoding two opposite body sides has high cognitive costs in younger children and smaller or null costs in older children. We interpreted these results in terms of the enhancement of Working Memory (WM) and body knowledge with age. When WM increases, it paradoxically drives older children to apply their superior body knowledge and imitate ‘anatomically’, producing side errors. Younger children are free from such interference because they still lack the necessary body knowledge and WM capacity. In conclusion, this study suggests that anatomical imitation becomes available in children’s cognitive repertoire from age nine due to increased body knowledge and WM capacity.

**Research Highlights:**

- The analysis of error types uncovers/illustrates the role of working memory and body knowledge in imitation.
- As an effect of little working memory capacity, younger children pay a cognitive cost to encode opposite body sides and make “side errors”.
- Imitation performance improves with age, but side errors paradoxically increase in frequency due to greater ability to master body knowledge bringing to anatomical imitation.
- Anatomical imitation is available in childrens’ cognitive repertoire from age 9, as an effect of the development of body knowledge and working memory capacity.

## Introduction

Imitation of others’ actions is crucial in acquiring new knowledge through social learning in infants and children. Human beings imitate their fellow humans to acknowledge several new abilities following their example, and such ability seems to originate early in life (Carpenter et al., 1998; Heyes, 2001; Meltzoff, 1996). Imitation guarantees the most straightforward learning mechanism, especially when compared to the less parsimonious “trials-an-errors” learning (Tomasello, 2003): observing someone’s actions allowed humans to learn faster and easier to relate with objects and environment over time, especially during early childhood (Meltzoff et al., 2009). Indeed, “imitation” refers to the ability to learn to perform an action and the ways to achieve it by reproducing both observed goals and gestures (Longo et al., 2008; Lyons et al., 2011; Want & Harris, 2002; Zentall, 2012). Thus, for “true” imitation to occur, both goals and the means to achieve must be reproduced, contrary to “emulation,” where only the goal is reproduced, and individuals acquire characteristics of objects and the environment without necessarily learning actions themselves. Importantly, these two processes seem to follow different developmental trajectories (Jones, 2009; McGuigan & Whiten, 2009; Want & Harris, 2002).

Imitation is acknowledged to rely on complex cognitive mechanisms involving observation, planning, and control of the action. For instance, the *Goal-directed theory* (GOADI; Bekkering et al., 2000) assumes that infants first need to decompose the observed actions in their constituent components, select some goal aspects, and then reconstruct them following a hierarchy among goals. However, in reproducing such goal hierarchy, they are influenced by cognitive resource constraints: if the action is too complex or new, only the main goals will be reproduced. GOADI assumes that the processes involved in imitation are exactly the same in both children and adults, with the only difference concerning working memory (WM) capacity (Wohlschläger et al., 2003). GOADI also states that the goal directly elicits the motor program most strongly associated with the seen action, highlighting the role of long-term memory. The *Dual-route model* (Cubelli et al., 2000; Rothi et al., 1991; Rumiati & Tessari, 2002) further develops on the existence of two independent processes of imitation: a direct route, used for imitating the new, meaningless actions but that can also be used for imitating known, meaningful ones when necessary, and a “lexical-semantic” (indirect) route which capitalizes on a long-term memory storage of known actions (the “lexicon”), and can only be used to imitate this kind of material (Rothi et al., 1991). Some authors also focused their attention on the role of WM in the processing of the direct route, as the action’s subunits need to be stored until they are recomposed in the motor output (Buxbaum & Randerath, 2018; Cubelli et al., 2000; Rumiati & Tessari, 2002; Tessari & Cubelli, 2014). Moreover, WM allows the two routes to interact in a shared working space to learn new actions (Tessari et al., 2006). Even though both routes are activated simultaneously (much like in other famous dual-route models, e.g., (Coltheart et al. ‘s DRC, 2001), the specific demands of an imitation task can modulate the emphasis on the information coming from a route or the other (Cubelli et al., 2006; Tessari & Rumiati, 2004; Tessari & Cubelli, 2014 for a critical discussion) and the available cognitive resources play an essential role (Tessari et al., 2006).

The study of imitation in children has always been of interest in the psychological as well as in the non-psychological literature (Hurley & Chater, 2005; Meltzoff, 1988; Want & Harris, 2002; Wohlschläger et al., 2003). Developmental studies on imitation have generally been interested in studying infants and preschool children to assess the presence of imitative mechanisms suggestive of acquisition of specific cognitive abilities (Dickerson et al., 2012; Huang & Charman, 2005; Nielsen, 2006). The question of *when* children and infants begin to imitate is still a matter of debate. Some authors propose that imitation is innate (Meltzoff & Moore, 1997, Nagy et al., 2005, Simpson et al., 2014), whereas others propose that it develops during the first years of life (Jones, 2007; Ray & Heyes, 2011). Some studies suggest that imitation develops with time: children become good imitators by 3LJyears of age (Horner and Whiten, 2005, McGuigan et al., 2007, Piaget, 1962), with automatic imitation quickening between 3 and 7 years (O’Sullivan et al., 2018).

### Purpose

So far, research on imitation in children generally focused on very simple actions and the role of movement complexity has been investigated in few studies. For example, Bauer (1992) demonstrated that children generally parse what they see in an appropriate way, but only few studies (e.g., Whiten et al., 2009) investigated the role of complex hierarchically structured action in imitation.. Also, imitation performance has been typically investigated with an eye on overall accuracy, with little attention to the pattern of errors produced while reproducing an action. Finally, the model that the child has to imitate is typically an adult. The purpose of the present work is to investigate imitation performance in children while bridging all these gaps.

By the mean of a cross-sectional approach, this study investigated how imitation develops through age in first-grade school children (6-10 years). We did not only focus on overall accuracy, but we decided to analyse different error types specifically. Indeed, the overall error rate is a composite measure reflecting several separate error categories, which can show different, even opposite and clashing effects. In the present context, error patterns are likely to be very informative on the role played by other cognitive steps and may change with age. Specific error rates might help understand how imitation develops and its relation to motor complexity and body representation development. Error analysis proved very fruitful in (neuro)psychology in general (Newcombe & Marshall, 1973; Toraldo & Shallice, 2004) and in the literature on imitation in adults, where it helped understanding which imitation process, direct or lexical route, was involved in performing the task (Carmo & Rumiati, 2009; Tessari & Rumiati, 2004) or was damaged in patients (Tessari et al., 2007).

We required children to imitate new, meaningless actions and decided to use actions with an increasing gradient of complexity: single limb movements (the simplest ones), two-limbs (one arm and one leg) homolateral movements, and two-limbs (one arm and one leg) contralateral movements (the most difficult ones) were used. Meaningless actions were chosen to prevent children from relying on previously acquired knowledge. Moreover, we used three models: a human child, a human adult and a child robot (iCub). Child and robot models might share the same body proportions and motor and spatiotemporal orientation faculties of the participants compared to the adult model. However, several studies (McGuigan et al., 2011; Rakoczy et al., 2010; Wood et al., 2012) have shown a bias in favour of adult models for imitation of novel or unusual, irrational or irrelevant actions.

## Method

### Participants

One hundred seventy-four children (88 females; mean age=8.25 years, sd = 1.34, range 6-11) participated in the experiment. All reported normal or corrected-to-normal vision and were nai[ve to the purpose of the experiment and were typically developing children (i.e., no cognitive delays, specific cognitive impairments, ADHD or specific learning disabilities). Fifty-seven children imitated the adult model (17 children were 6-7 years old; 17 were 8 years old; 11 were 9 years old and 12 were 10-11 years old); sixty-two children imitated the action performed by the peer model (15 children were 6-7 years old; 15 were 8 years old; 19 were 9 years old and 13 were 10-11 years old); and fifty-five children imitated the action performed by the robot model (17 children were 6-7 years old; 15 were 8 years old; 11 were 9 years old and 12 were 10-11 years old).

### Ethical Approval

Ethical approval was granted by the Bioethics Committee of the University of Bologna on the 7^th^ of May 2013, following the Declaration of Helsinki, and parents/tutors provided written informed consent for the children to participate in the study.

### Stimuli

Children were shown three different models: a 10-year-old child, a 22-year-old boy, or an iCub robot. They performed 12 meaningless actions: 4 single effector gestures involving only one leg or one arm (“Single” condition); 4 ipsilateral movements involving one arm and one leg belonging to the same hemisome (“Homolateral” condition); 4 contralateral movements involving one arm and one leg belonging to two different hemisomes (“Heterolateral” condition; they were matched with the movements in the homolateral condition). The action videos were flipped along the horizontal axis to generate the mirror image of each action, for a total number of 24 stimuli. The iCub videos were created using the humanoid platform simulator iCubSim (Tikhanoff et al., 2008). See Figure 1 in Ottoboni et al. (2013) for some examples.

**Figure 1:**
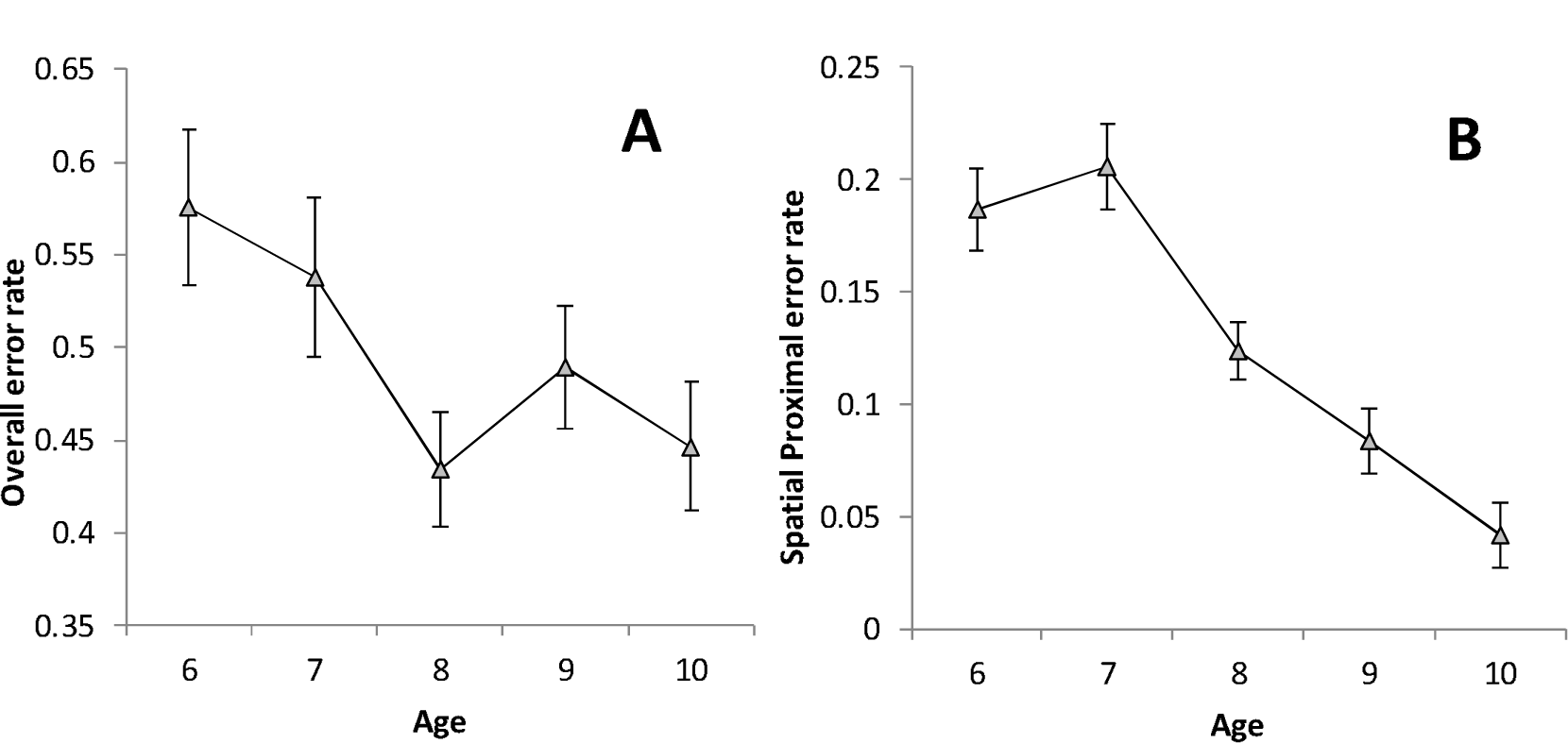
Overall error rate (A) and Spatial Proximal error rate (B) as a function of Age (Mean ± Standard Error).

### Procedure

Participants kept the spacebar pressed at the beginning of each trial while observing the model’s action. A red circle (Go signal) informed the participants that they could release the spacebar and imitate the just observed action as if they were in front of a mirror (we chose specular imitation as it is more natural in children, Brass et al., 2000). Once imitation was concluded, the participants were required to press the spacebar again, and the subsequent trial began. All videos were projected from 3 meters away from the children (using a DELL 2300 projector), who were encouraged and supported after the imitation, irrespective of the quality of the gesture. Imitation was video-recorded and later scored by two independent raters (Coher’s Kappa = .86). They reported and extracted all the error types found in the videos and later discussed their typology with the other authors (G.O, A.To, and A.Te). On the bases of these detailed analyses, 10 different types of error emerged:

- *Spatial distal*: hand or foot moved incorrectly or not moved at all, while the involved limb was moved correctly.
- *Spatial proximal*: limb moved with a wrong angle with respect to the trunk or limb moved correctly on the wrong orientation plan.
- *Spatial final position*: wrong final position of the involved limb.
- *Perseveration*: action performed is completely different from the target but is ascribable to another action previously observed (global perseveration) or action performed is partially correct and partially ascribable to another one previously observed (partial perseveration).
- *Omission* an action step that is necessary for completing the movement is not carried out.
- *Sequence:* a synchronous action of the arm and the leg has been temporarily decomposed (the arm anticipates the leg or vice versa).
- *Side*: it occurs when imitation is anatomical and not mirror-reversed for the arm, leg, or both.
- *Conduit d’approche:* the children attempt to correct their mistakes through spontaneous corrections, with gestures progressively coming closer and closer to the target.
- *Arm/Leg:* an observed arm movement is carried out with the leg and vice versa.
- *Unclassifiable:* the child reproduced a completely different gesture compared to the presented one, not due to one of the previous errors.

An imitated action was scored wrong when at least on one of these errors occurred.

### Statistical analyses

Generalized Mixed Linear Models (GMLM) with Binomial distribution and logit link were used to analyse error rates. The models included the full factorial design Condition (Single effector vs. two Homolateral effectors vs. two Heterolateral effectors) x Model (Adult vs. Child vs. Robot) x Age (categorical). Given that Condition was the only within-subjects factor, the random design contained the by-subject random intercept and the Condition random slope. When Age effects were significant and theoretically interesting, we tried to understand at what age it “flattens” / stabilizes (when it is a decreasing curve) or it begins to rise (when it is an increasing curve). To this purpose, non-linear regressions (*nls* in R) were run on the per-subject error rates (mean across the three Conditions). Two models were considered, which took into account the bounded nature of the error rate scale, a logistic model with equation Error Rate = *fl* + *vr*/(1+exp(ln((1-0.05)/0.05)+*w*\**s*-*s*\*Age)), and (only for decreasing patterns) an exponential decay with equation Error Rate = *fl* + *vr*\*exp((Age-6)*ln(0.05)/*w*). In both cases, performance is thought to have a lower asymptote (*fl*, for “floor”), and a “vertical range” (*vr*) in which it varies, with slope *s*; *w* is the age when performance stabilizes or begins to rise, and is defined as the point where the function is 5% (hence the 0.05 in the equations) of the vertical range above the lower asymptote (see Appendix A for details).

Given that we analysed the overall error rate as well as nine specific error rates (those related to the nine error categories), a Bonferroni correction was applied in the latter set: effects whose p < .05/9 = .0055 were considered to be significant. General Equation Estimation (GEE) was used to test specific hypotheses on the error categories in which the Heterolateral condition yielded higher error rates than the Single-effector condition. Those hypotheses required data transformations that prevented the use of GMLM, but still allowed GEE to properly model the effect of a repeated-measure variable (Condition) on error rates characterized by very irregular (Tweedie) shapes.

We were unaware of any previous experiment with a similar method and population, so we lacked empirical effect size estimates. Therefore, we ran a sensitivity analysis, which provided the minimal detectable effects that the present experiment can grant with power fixed at conventional 0.8 and sample size fixed at the (actual) *N*=174. GMLM results were approximated using a classical GLM approach (Gpower 3.1.9.4, Faul et al., 2007); such approximation should, if anything, underestimate sensitivity given that GMLM is generally more powerful than GLM (Baayen, 2008). Table 1 lists the detectable effect sizes, ranging between 0.09 and 0.23 SD (small-medium range, Cohen, 1969, p. 348), except Model x Age interaction, which ranged from 0.3-0.37 (medium-large).

**Table 1:** Detectable effect sizes (Cohen’s *f*) from Sensitivity Analysis

| **Table 1:** Detectable effect sizes (Cohen's *f*) from Sensitivity Analysis |  |  |
| --- | --- | --- |
|  | Overall Error Rate | Specific Error Rates |
| Condition (within) | .0966 | .1502 |
| Model (between) | .1937 | .2133 |
| Age (between) | .2167 | .2349 |
| Condition*Model | .1078 | .1648 |
| Condition*Age | .1215 | .1833 |
| Model*Age | .301 | .3726 |
| According to Cohen's convention (1969, p. 348) $f = .1$ is "small", $f = .25$ is "medium" and $f = .4$ is "large". Input to Gpower 3.1.9.4 was: power = 0.8; $N=174$; alpha = 0.05 for Overall Error rate and .0055 (Bonferroni-corrected) for Specific Error Rates; average correlations were set at (empirically-derived) average values of .4985 and .2476 for Overall and specific rates respectively. Gpower 3.1.9.4 does not provide sensitivity analysis for triple (between-within) interactions. |  |  |

## Results

Table 2 reports details about the types of errors occurred in the imitation task ither singularly or in combination.

**Table 2:** Report of error types

| **Table 2: Report of error types** |  |  |  |
| --- | --- | --- | --- |
|  | Raw n. of error | % out of all errors | % out of all trials |
| SPATIAL DISTAL | 220 | 10.87 | 5.27 |
| SPATIAL PROXIMAL | 486 | 24.01 | 11.64 |
| SPATIAL FINAL | 218 | 10.77 | 5.22 |
| PERSEV | 158 | 7.81 | 3.78 |
| OMISS | 50 | 2.47 | 1.20 |
| SEQ | 217 | 10.72 | 5.20 |
| SIDE | 604 | 29.84 | 14.46 |
| CONDUIT APPR | 118 | 5.83 | 2.83 |
| ARMLEG | 186 | 9.19 | 4.45 |
| Unclassifiable | 50 | 2.47 | 1.20 |
| Double classifications | -271 | -13.39 | -6.49 |
| Triple classifications<sup>1</sup> | -12 | -0.59 | -0.29 |
| Overall | 2024 | 100.00 | 48.75 |

Due to the number of factors and interactions and to facilitate understanding of the results from GMLM analyses, they are shown in Table 3.

**Table 3:** Results from Generalized Mixed Linear Models

|  | *Overall*<br>*Error*<br>*rate*<br>*(4131)* | *Spatial* |  |  | *Persev*<br>*(4131)* | *Omiss*<br>*(4131)* | *Seq*<br>*(4131)* | *Side*<br>*(4147)* | *Con*<br>*Appr*<br>*(4131)* | *Armleg*<br>*(2754)\** |
| --- | --- | --- | --- | --- | --- | --- | --- | --- | --- | --- |
| *Predictor*<br>*(df)* |  | *Distal*<br>*(4131)* | *Proximal*<br>*(4147)* | *Final*<br>*(4131)* |  |  |  |  |  |  |
| *Age (4)* | **2.609,**<br>**.034** | 1.003,<br>.405 | **11.566,**<br>**<.001** | .534,<br>.711 | .48,<br>.751 | .121,<br>.975 | 2.867,<br>.022 | **9.864,**<br>**<.001** | 1.409,<br>.228 | .674, .61 |
| *Condition*<br>*(2)\** | **34.241,**<br>**<.001** | **76.627,**<br>**<.001** | **40.543,**<br>**<.001** | **14.321,**<br>**<.001** | .213,<br>.808 | .42,<br>.657 | 4.931,<br>.007 | **58.28,**<br>**<.001** | 3.314,<br>.036 | 3.444,<br>.064 |
| *Model (2)* | .463,<br>.629 | .699,<br>.497 | .907, .404 | 4.412,<br>.012 | 1.249,<br>.287 | .5, .607 | 1.996,<br>.136 | 4.928,<br>.007 | 1.636,<br>.195 | .268,<br>.765 |
| *AxC (8)\** | 1.115,<br>.35 | 1.53,<br>.141 | 1.435,<br>.176 | .437, .9 | 1.33,<br>.223 | .031, 1 | 1.097,<br>.362 | 2.255,<br>.021 | .358,<br>.943 | .3, .878 |
| *AxM (8)* | 1.781,<br>.076 | .098,<br>.999 | 1.85, .063 | .999,<br>.434 | .379,<br>.932 | .237,<br>.984 | .443,<br>.895 | 1.118,<br>.347 | .335,<br>.953 | .89, .524 |
| *CxM (4)\** | **7.631,**<br>**<.001** | .901,<br>.462 | .927, .447 | .645, .63 | 1.001,<br>.406 | .182,<br>.948 | **6.448,**<br>**<.001** | 1.507,<br>.197 | .124,<br>.974 | .227,<br>.797 |
| *AxCxM*<br>*\*(16)* | 0.896,<br>.574 | .184, 1 |  | .397,<br>.984 | .403,<br>.982 | .116, 1 | .544,<br>.925 |  | .137, 1 | .351,<br>.946 |

Each cell reports F and p-values, which are in **bold** when significant (Bonferroni correction applied to specific error categories). Different rows report effect by different predictors. Interactions contained abbreviated variables names, A=Age (analyzed as a categorical variable), C=Condition, M=Model. Different columns report different error categories (dependent variables). Values in brackets indicate degrees of freedom. \*Armleg: single-effector trials were omitted, as no arm-leg confusion errors could have occurred in this case; thus numerator's degrees of freedom were 1 for Condition, 4 for AxC, 2 for CxM, 8 for AxCxM. Empty cells: triple interactions were omitted due to convergence failures.

### Overall error rate

The GMLM analysis detected a significant effect of Age (F(4, 4131)=2.609, p=.034): error rate showed some moderate drop with Age (Figure 1A). The logistic model did not converge, so we applied the exponential decay model that gave the following parameter estimates (mean±standard error): *floor*=.453±.035; *vertical range*=0.129±.051; *stabilization Age* = 8.97±3.24 years. Thus, performance flattened at about age 9, but this parameter has a very unstable estimate (a standard error over 3 years).

Condition yielded a significant effect (F(2, 4131)=32.241, p<.001), but with a strong Condition x Model interaction (F(4, 4131)=7.631, p<.001) shown in Figure 2A. Indeed, Single-effector gestures were the easiest, Heterolateral gestures were the most difficult, and Homolateral had intermediate difficulty. However, this only happened when the model was either a Child (F(2, 1473) = 22.808, p<.001) or a Robot (F(2, 1305) = 21.838, p<.001): when the model was an Adult, such effects disappeared (F(2, 1353) = 1.331, p=.264). Model had a significant effect in the most difficult, Heterolateral condition, where the Adult model showed a sizeable, 15.2% advantage over Robot and Child models (F(2, 1377)=4.762, p=.009), while it did not reach significance in the Homolateral condition (F(2, 1377)=1.316, p=.269) and in the Single effector condition (F(2, 1377)=2.53, p=.08), where the effect was in the opposite direction (children performing worst with the Adult model, Figure 3A).

**Figure 2:**
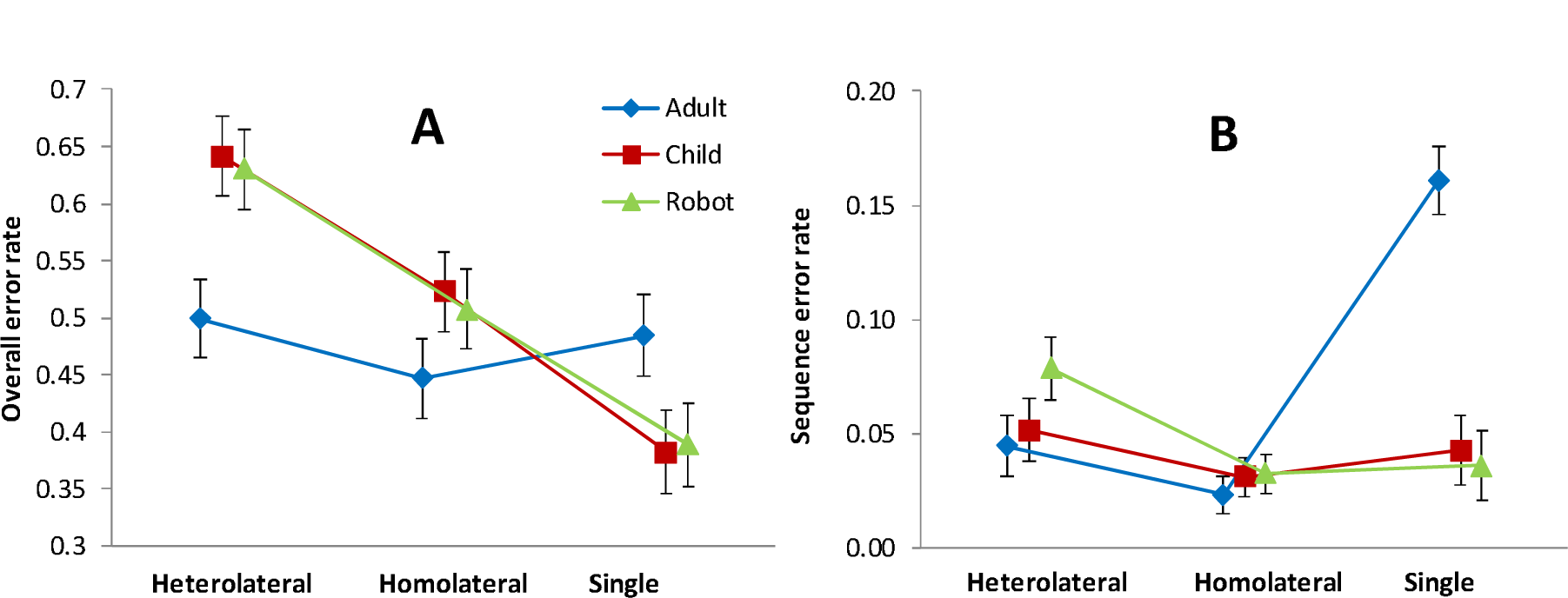
Condition x Model interaction (Mean ± SE) on Overall error rate (A) and Sequence error rate (B).

**Figure 3:**
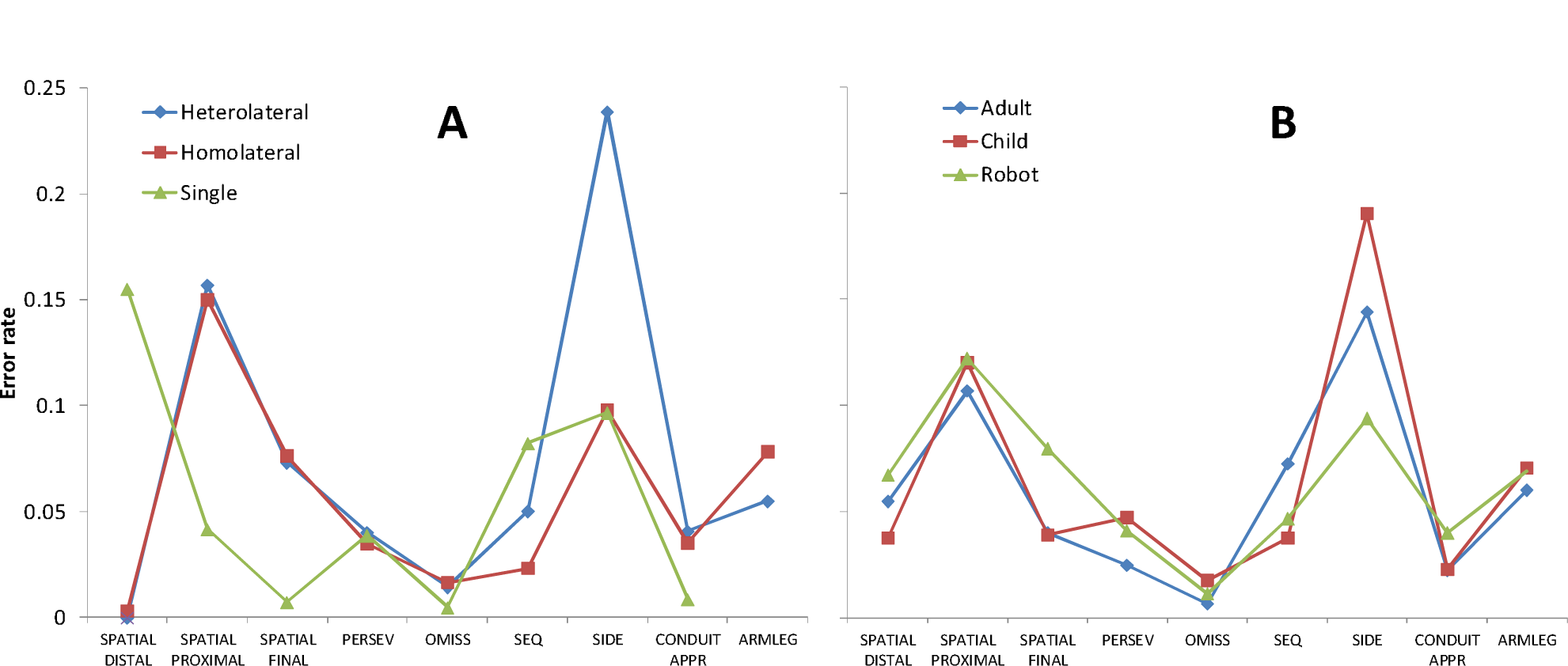
Error rate for specific error categories as a function of (A) Condition, and (B) Model. In A, Are-leg confusion errors were impossible if a Single effector was involved.

Importantly, the overall error rate is a composite measure reflecting several separate error categories, which show different, even opposite and clashing effects (see Figures 4A and 4B), so a straightforward interpretation is difficult. Analyses of specific error categories will clarify this issue.

#### Error profiles

To provide an overview, Figure 3 shows error rates for specific error categories as a function of Condition (Figure 3A) or Model (Figure 3B). We could not perform statistical comparisons *across* error categories as these are not mutually independent (they are subsets of a constant overall set of trials, so their frequency values constrain each other). Only qualitative comparisons are allowed in this case. As to within-error-category comparisons, statistical tests are reported later.

Figure 3 shows that, in general, Spatial Proximal and Side errors were the most common ones. Interesting within-category differences, according to either Condition (Figure 3A) or Model (Figure 3B) emerged, which are analyzed below.

#### Statistical analysis within specific error categories

Perseveration, Omission, Conduit d’approche, Arm-leg (fusion or inversion) error rates failed to show (Bonferroni-corrected) significant effects (Table 2), so we focused on other error categories.

### Side errors

The triple interaction had to be removed from GMLM due to convergence failure. The analysis yielded massive effects of Condition (F(1, 4147) = 58.28, p<.001) and Age (F(4, 4147) = 9.864, p<.001). The rate of Side errors clearly *increased* with Age (Figure 4). Nonlinear regression (logistic model) delivered the following estimates: *floor* = .068±.027; *vertical range* = 0.195±.05; *slope* = 2.83±2.15; *rising Age* = 7.72±0.89. Thus, at about Age = 8, error rate begins to rise.

Concerning the effect of Condition (visible in Figure 3A), as expected, Side errors were most frequent when spatial and bodily encoding demands are highest, i.e., in the Heterolateral condition. In the Homolateral and Single-effector conditions error rates were much lower, without any sizeable differences between them. Thus, when a single side of the body was involved, subjects encoded its side relatively easily, no matter whether the gesture recruited one or two effectors. By contrast, spatial encoding was much less accurate when both body sides were involved (in a crossed pattern).

Albeit just liminal due to the Bonferroni correction (F(2, 4147)=4.928, p=.007>.0055), the effect of Model on Side errors is worth mentioning, insofar as it involves very large (two-fold) differences in relative frequency: when the model was a child, Side errors were 19%, while a robot only induced 9% such errors (adult model: 14%, Figure 3B). This pattern was confirmed and investigated in more depth in the following GEE analysis.

### General Equation Estimation (GEE) to test the “cumulative probability” hypothesis

Side errors were markedly more frequent in the Heterolateral than in the Homolateral and Single-effector conditions (Figure 4). The most straightforward explanation for such a difference is that an error is more likely to occur if one has more “opportunities” to commit one: if one has to encode two sides – the side of the arm and the side of the leg, as it happens with Heterolateral actions, the probability of making at least one mistake is higher than when only one side is to be encoded, as in Single-effector actions, because of sheer combinatorics. The question then is whether the disadvantage shown by Heterolateral actions was *entirely due* to such a “cumulative probability” phenomenon or whether there were extra difficulties that those actions imply. Luckily, we can try to answer this question because the cumulative probability is a mathematically known factor that can be “removed” from the Heterolateral error rate. If cumulative probability were the only reason for the Heterolateral disadvantage, nothing of that disadvantage would remain after the mathematical removal, and the adjusted Heterolateral score would match that of the Single-effector score. Else, if there were other, genuinely cognitive, reasons that make the Heterolateral condition more difficult, after the adjustment, the Heterolateral error rate would remain higher than the Single-effector one.

The mathematical adjustment worked as follows. If cumulative probability were the only factor at play, and *x* were the Side error rate in the Single effector condition, the expected Side error rate in the Heterolateral condition would be *y*=2*x*-*x*^2^. Thus, the inverse transformation, *x* = 1-(1-*y*)^1/2^, applied to the Heterolateral error rate would provide us with a valid estimate of the Single-effector error rate, to be compared to the observed one (see Appendix B for details). If the two match each other, cumulative probability explains all the Heterolateral disadvantages; otherwise, the Heterolateral condition is inferred to bear some extra cognitive difficulties.

Adjusted Heterolateral scores were obtained from each subject and compared to the average error rates from the Single and Homolateral conditions (which proved virtually identical, see previous GMLM and Figure 4, thus averaging further increased statistical power). The comparison was carried out by a GEE analysis (previously validated in a small Monte Carlo study) with Age (five classes) and Model (three types) as between-subjects factors, and Condition as within-subjects factor (Adjusted Heterolateral vs Single/Homolateral Side error rates)^1^. Distributions of Side error rates were modelled as Tweedie with log link. Table 4 reports GEE results.

**Table 4:** Results of GEE on Side error rates

|  | *Wald Chi-square* | *Df* | *p* |
| --- | --- | --- | --- |
| Age | 50.564 | 4 | <.001 |
| Model | 7.539 | 2 | .023 |
| Condition | 38.299 | 1 | <.001 |
| Age x Model | 8.417 | 8 | .394 |
| Age x Condition | 25.332 | 4 | <.001 |
| Model x Condition | 7.713 | 2 | .021 |
| Age x Model x Condition | 19.056 | 8 | .015 |

Results were very interesting and clear-cut. Condition had a massive effect and strongly interacted with Age and Model, thus definitely rejecting the hypothesis that the additional difficulty of the Heterolateral with respect to the other conditions was only due to the cumulative probability of a Side error across limbs. Clearly, there are extra-difficulties, which are well visible as the gaps between the adjusted Heterolateral (blue) and the Single/Homolateral (red) scores in Figure 5A. The Condition x Age interaction (Table 3, Figure 5A) showed that extra-difficulties characterized the performance of younger children (6-8 years: Wald χ^2^= 35.081, p<.001) and much less so that of older (9-10) children (Wald χ^2^=3.281, p=.07), whose pattern of results was compatible with the “cumulative probability” hypothesis (their post-adjustment probability of a Side error was similar in both conditions, right side of Figure 5A). The other interesting evidence came from the interactions involving Model, especially from the triple interaction (Table 3). This was due to the fact that the Model x Condition interaction was undetectable among older (9-10) children (Wald χ^2^=.665, p=.717), but significant among younger (6-8) ones (Wald χ^2^=9.112, p=.011); in turn, the latter interaction arose because the Condition effect was very strong with a Child model (Wald χ^2^= 36.962, p<.001), weaker but still significant with an Adult model (Wald χ^2^= 9.29, p=.002), and not significant with a Robot model (Wald χ^2^= 2.677, p=.102). Figure 5B clarifies the pattern of the triple interaction, by showing the Condition effect on the Y axis: hence, the axis directly illustrates the presence and intensity of extra-difficulties in Heterolateral gestures that are not merely due to the number of sides (two) to be encoded. Such extra-difficulties are generally very small and close to zero, except for younger children watching a Child model.

**Figure 5:**
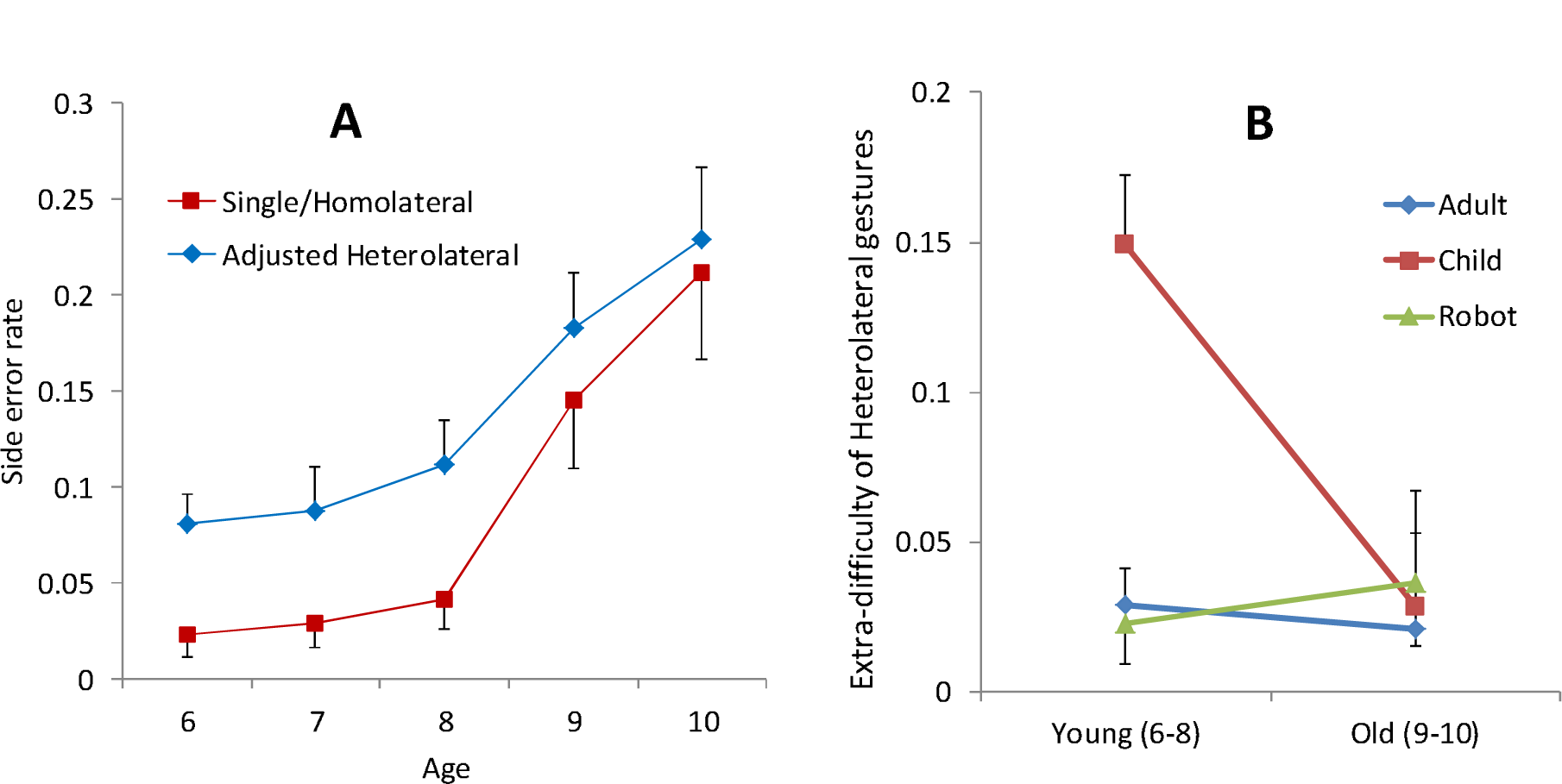
A. Side error rate (Mean ± SE) as a function of Age (years). Red: average between Single and Homolateral error rates (they showed no difference in previous GMLM analyses); blue: Adjusted Heterolateral error rates (i.e. after removal of the cumulative-probability factor). B. Extra-difficulty of Heterolateral gestures with respect to other gestures (Y axis) is plotted as a function of Age (merged classes: 6-8 yrs, 9-10 yrs, X axis) and Model (blue, Adult; red, Child; green, Robot). Extra-difficulty (Y) was computed as the difference between the Adjusted rate of Side errors in the Heterolateral condition and the average rate of Side errors in the Single and Homolateral conditions (see text for details). Table 3 reports GEE analyses’ significance levels.

Wrapping up, older children (9-10) had much *higher* Side error rates than younger (6-8) ones. Yet, they did *not* show extra-difficulties in the Heterolateral condition, over and above the mere cumulative probability of making an error on any limb (Figure 5A). By contrast, younger children (6-8) made much *fewer* Side errors; however, when the model was a Child, they showed marked extra-difficulties in the Heterolateral condition, which cannot be explained away by the cumulative probability of an error (Figures 6A, 6B).

**Figure 6:**
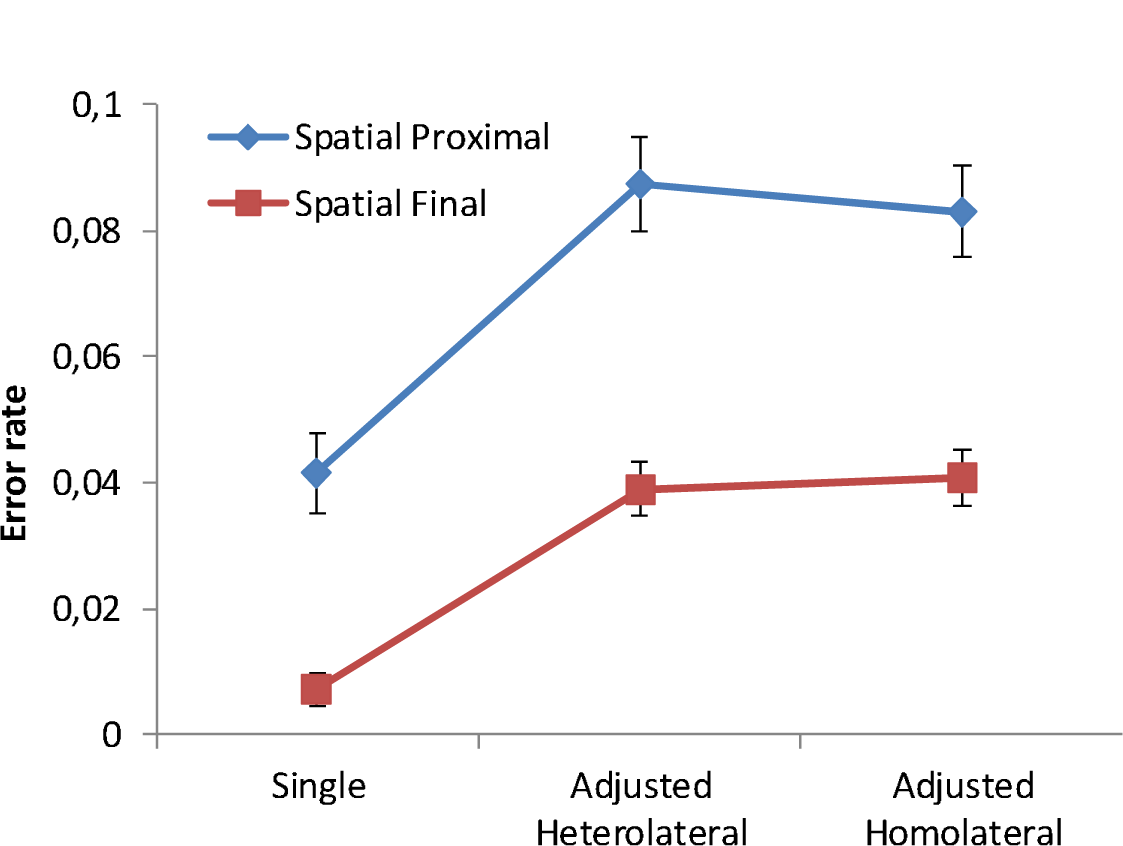
Error rates as a function of Condition; Heterolateral and Homolateral rates were adjusted to remove the cumulative-probability factor; blue, Spatial Proximal; red, Spatial Final.

#### Spatial distal errors

Condition had a massive effect (F(2, 1431)=76.627, p<.001). As visible in Figure 3A, the percentage of distal spatial errors was virtually nihil with double-effector (both Heterolateral and Homolateral) stimuli and much higher (15.4%) in Single movement stimuli. This apparently paradoxical result is actually trivial: “single-effector” stimuli contained distal components that were not included in double-effector stimuli.

#### Spatial Proximal errors

The full model did not converge on reliable solutions (yielding very high standard errors of parameter estimates), so we excluded the triple interaction from the model. Results are shown in Figure 3A. Single-effector gestures produced *fewer* Spatial Proximal errors than double-effector ones (Condition: F(2, 4147)=40.543), p<.001). Moreover, a massive effect of Age emerged (F(4, 4147) = 11.566, p<.001), with error rate being lower in older participants. Nonlinear regression (logistic model) gave estimates: *floor* = .044±.023; *vertical range* = 0.158±.038; *slope* = −1.81±1.06; *stabilization Age* = 9.75±1.02. In this case, the logistic fit identified a stabilization point which essentially corresponds to the oldest age class tested, because those participants had very close-to-zero error rates (Figure 1B).

Like Side errors, also Spatial Proximal errors showed a strong disadvantage for Heterolateral with respect to Single gestures, so the question of whether this is a mere effect of cumulative probability of an error over the two limbs, or also the effect of extra-difficulties, came to the fore. Thus, a GEE analysis was applied to Spatial Proximal errors; however, this time the Homolateral condition behaved as the Heterolateral one (Figure 3), so both underwent the 1-(1-error rate)^1/2^ transformation at the level of single subject to partial put the cumulative-probability factor. GEE yielded a massive Condition effect (Wald χ^2^=27.679, p<.001), with both the Heterolateral (Wald χ^2^=28.768, p<.001) and Homolateral (Wald χ^2^=21.914, p<.001) conditions showing a residual disadvantage with respect to Single, and without any significant difference between Heterolateral and Homolateral themselves (Wald χ^2^=.443, p=.506). Thus, the hypothesis of mere cumulative probability of an error was rejected, for both the Heterolateral and the Homolateral conditions (Figure 6). This held for both 6-7 year olds (Condition effect: Wald χ^2^=18.878, p<.001) and for 8-10 year olds (Wald χ^2^=13.953, p<.001). While Condition x Age did not reach significance (Wald χ^2^=15.361, p=.053), the absolute sizes of the Hetero/Homo vs Single gaps in performance were quite different (.09 for younger, .026 for older children), an expected pattern given that the performance of older children was much closer to the floor.

#### Spatial Final-Position errors

Condition (F(2, 4131) = 14.321, p<.001) yielded a significant effect, due to the usual advantage of Single effector (about 3% error rate) over the other conditions (both at about 8% error rate), visible in Figure 3A. Such a pattern called for another GEE analysis, to understand whether or not a simple cumulative-probability pattern characterized the processing of double-effector gestures. Again, the 1-(1-error rate)^1/2^ transformation was applied to obtain adjusted error rates from both Heterolateral and Homolateral raw rates given that they were both higher than the Single raw rate (Figure 3A). Like with Spatial Proximal errors, the Spatial Final error rate (analyzed via GEE) showed a strong Condition effect (Wald χ^2^=23.531, p<.001), due to clear-cut advantage of Single over both Adjusted Heterolateral (Wald χ^2^=25.879, p<.001) and Adjusted Homolateral (Wald χ^2^=24.722, p<.001), and without differences between the latter two (Wald χ^2^=.427, p=.513). Again, the simple cumulative probability hypothesis was rejected, with both Heterolateral and Homolateral conditions posing some extra-difficulties to participants (Figure 6). This held for both 6–7-year-old (Condition effect: Wald χ^2^=7.458, p=.024) and for 8-10 year olds (Wald χ^2^=19.097, p<.001), with virtually identical absolute sizes (error rate differences between Homo/Hetero and Single were .036 and .031 respectively^2^).

#### Sequence errors

A highly significant Condition x Model interaction (F(4, 4131) = 6.448, p<.001) was found, which was clearly due to *adult-model, single gestures* inducing definitely more sequence errors than all other combinations (Figure 2B). Indeed, Model had a marked effect in the Single condition (F(2, 1377) = 16.902, p<.001) while it fell short of significance in the Homolateral (F(2, 1377) = .137, p = .872) and in the Heterolateral (F(2, 1377) = .98, p = .376) conditions.

## General discussion

This study investigated the developmental trajectory of imitation in first-grade school children (aged 6-11years old). We manipulated the complexity of meaningless actions (from movements of a single limb to movements involving arm and leg of the same body side (Homolateral) to movements involving arm and leg of opposite sides (Heterolateral), and used three models (a human child, a human adult and a child robot, the iCub). We found several interesting and clear-cut patterns, each of which deserves detailed discussion, which we provide in the following paragraphs.

### Role of model

Previous literature suggested that an adult model acquires a role of behavioural guidance on children’s imitation through their supposed authoritativeness: when *new* actions are shown, children will tend to rely more on adults than on peers in imitation as they tend to acquire information from adults for unfamiliar domains but equally often from peers and adults for familiar domains (Taylor et al., 1991). Children infer the expertise and the reliability of the model based on the inferred expertise on specific domains (Chow et al., 2008; Jaswal & Malone, 2007; Jaswal & Neely, 2006), and adults are considered in general more reliable in light of their role for social learning (Laland, 2004). Given that we used meaningless, novel actions as stimuli in our study, the studies mentioned above would predict that an adult model should yield a better imitation performance than a peer or a robot one. When analysing overall error rate, this prediction was only partially confirmed, as such a pattern characterized only the most difficult, Heterolateral gestures, and to a lesser extent, Homolateral ones – the simplest gestures, involving a single limb, showed, if anything, a paradoxical disadvantage of adult-model gestures (Figure 2A), possibly related to Sequence errors (Figure 2B). We have no straightforward explanation for the latter result. In the context of Side errors, we did find an advantage for both the adult and the robot model over the child model in younger (6-8 years old) children for Heterolateral gestures (Figure 5B). One possibility is that, for different reasons, the adult and the robot model increase young children’s attention, the adult model for the authoritativeness effect referred to above, and the robot model by involving children in a game-like interaction (as showed in a wide range of studies on autistic children, e.g., (Michaud et al., 2003; Michaud & Théberge-Turmel, 2002) or in healthy participants in a physiotherapy session; (Brooks & Howard, 2012)

### Role of Age

A second interesting finding regards the effects of Age. While overall performance improves with age (Figure 1A), and error rates stabilize at about 45% at 9 years (a result that is confirmed in some error categories, especially Spatial Proximal, Figure 1B), exactly the opposite holds true for Side errors (Figure 4). These mistakes are stable in frequency (and virtually absent in two sub-categories) at ages 6 to 8, then their frequency suddenly rises at age 9, and increases even further in 10-year-olds. We believe the most plausible explanation for such an unexpected result is that older children might experience some form of interference by an “anatomically”-based imitation strategy, which might simply be absent from the cognitive repertoire of younger children (see later), who would thus show the paradoxical, observed advantage. Thus, older children might sometimes fail to apply the instructions, which required them to imitate as if they were in front of a mirror, and imitate “anatomically”, that is, moving (e.g.) their right limb if the model moved their right limb.

### More on Side errors

The above explanation should be extended to account for other evidence obtained from Side errors. These were most frequent in the Heterolateral condition, which is hardly surprising given that Heterolateral are supposed to be the most difficult items. However, such a disadvantage was not due to the trivial fact that two limbs had to be moved rather than one, but to the fact that those two limbs were in a crossed pattern – left arm plus right leg, or vice versa. Indeed, Homolateral gestures also involved two limbs, which, however, were on the same side of the body, and Side error rate for them was as low as in the baseline condition that involved a single limb. We believe such a pattern can be interpreted by reasoning on the number of spatial representations (codes) that children internally produce to specify the laterality of the action. Clearly, the more codes are produced, the more likely it is that at least one of them is wrong (i.e., the higher the probability of a Side error to occur). We started from the hypothesis that children produced separate laterality codes for arm and leg. If this had been the case, double-effector movements would have shown more Side errors than Single-effector movements. Thus, for instance, a Homolateral trial involving the left arm and the left leg would be encoded as “arm=left” and “leg=left”, and errors would be more frequent than in the baseline condition involving a single code, e.g. “arm=left”. Facts contradicted this prediction: the Homolateral condition yielded a virtually identical Side error rate to that of the Single-effector condition – it was only the Heterolateral condition that yielded a (massively) higher Side error rate.

We believe the most plausible explanation is that in both the Single-effector and Homolateral-effectors conditions, children produced only one laterality code attached either to the single limb involved (Single condition), or to the arm-leg pair (Homolateral condition), e.g., “arm+leg=left”, thus having the same probability of making a Side error in both cases. By contrast, in the Heterolateral condition they could not help but encode two sides, the side of the arm and the side of the leg (e.g. “arm=left”, “leg=right”), and by using two codes, error probability unavoidably increased.

However, another question comes to the fore. Did the error rate increase *only* because moving two (crossed) limbs entails two spatial codes instead of one spatial code? Else, do Heterolateral gestures yield some extra difficulty to the cognitive system, over and above the mere cumulative probability of an error? We answered this question by mathematically removing the cumulative-probability factor from the error rates of the Heterolateral condition and looking at whether some residual disadvantage remains. Older children (9-10) showed no residual disadvantage: therefore, to them, Heterolateral gestures were more difficult just because they had two “opportunities” to make a Side error (the two limbs); by contrast, younger children (6-8) did show a residual disadvantage for Heterolateral gestures, thus suggesting the presence of some extra-source of difficulty in encoding two sides (Figure 5A). We believe such an extra difficulty might be due to either Short-Term Memory (STM) capacity, which is definitely smaller in younger children (Gathercole, 1999; Isaacs & Vargha-Khadem, 1989; Wilson et al., 1987), or to interference between the two codes: for instance, the “left” code for the arm might interfere with the “right” code for the leg, with younger children being less able to manage conflicting information. In any case, such limitations would be absent in older, 9-10-year-old children (whose Side error rates, after adjustment, were very similar in both the Heterolateral and Single/Homolateral conditions, Figure 5A). Older children did produce more Side errors than younger children – as hypothesized above, an effect of the appearance of an inadequate anatomical imitation strategy at about age 9 – but in doing so, older children did not show extra-difficulties due to STM limits or inability to manage conflicting information (these latter were specific of younger children).

In summary, Side errors showed a “double dissociation” between younger and older children (well visible in Figure 5A). Younger children have STM or conflict-managing limits, which produce extra-difficulty in encoding two sides instead of one, but are not affected by an anatomical imitation strategy, which is likely to be still absent from their cognitive repertoire. Older children have precisely the opposite pattern: they do not show extra-difficulties, as maturation provided them with sufficient STM and conflict-managing abilities for the material used in the present experiment, but sometimes apply an anatomical imitation strategy, which worsens their side encoding performance (see Appendix B for further reasoning on this point). Interestingly, the extra-difficulty experienced by younger children in the Heterolateral conditions almost only affected trials in which the model was a child (Figure 5B). We already discussed the possibility that the adult and the robot models elicit more attention/interest in young children. In this vein, a greater attentional involvement might mask the limited STM capacity and/or the limits in conflict-managing abilities, which in turn would emerge with a less engaging, child model.

### Spatial Proximal and Spatial Final-Position errors

The comparison between the pattern of Side errors, discussed above, and those of Spatial Proximal and Spatial-Final-Position errors is also interesting. In the latter two error types, we assessed how accurately the *content* – and not the side – of the movement had been processed. The Heterolateral condition yielded many more errors than the baseline, Single-limb condition (Figure 3A) also for the Spatial Proximal and Spatial Final categories. However, this time, as expected, just the *number* of limbs counted, and not their side: indeed, the Homolateral condition yielded virtually identical error rates as those of the Heterolateral condition. While the *side* of (e.g.) the left arm and the left leg could well be “merged” into a single spatial code (e.g. “left limbs”), thus making the Side error rate drop for the Homolateral condition, this was, of course, impossible when the content of the movement had to be encoded, because this was always different between arm and leg. Hence when looking at Spatial Proximal and Final error rates, which express the adequacy of the reproduction of the movement content, Heterolateral and Homolateral gestures were equally difficult.

The pattern related to putative STM and/or conflict-managing capacities was also different with respect to what we found with Side errors. With Spatial Proximal and Final errors, both younger and older children showed significant extra-difficulties when dealing with two limbs instead of one. Indeed, in both conditions with two effectors (Heterolateral and Homolateral), younger as well as older children had a higher error rate than that recorded in the Single-effector condition even after adjusting for the cumulative-probability factor (Figure 6).

Wrapping up, when the content of the gesture was concerned, all children, irrespective of age, showed a disadvantage for gestures involving two limbs with respect to those that involved only one limb, a performance gap that was not merely due to combinatorial probability, that is, to the fact that two limbs are more likely to produce at least one error than a single limb. STM limitation and/or conflict managing difficulties are, again, plausible explanations for such a pattern.

### Separate STM systems

It is critical to note that different STM systems are likely to be involved in storing different features of the action to be imitated. We have direct proof of such a notion in our data, regarding the *side* of the action versus the *content* of the action. By playing the devil’s advocate, suppose that a single STM system had been involved in storing both the side(s) of the effectors involved and the action’s content, that is, the exact movements to be reproduced. If this had been the case, Side errors should have been more frequent in the Homolateral than in the Single condition, because two different movements vs one movement would be involved respectively, thus leading to different STM loads. Another prediction is that the Heterolateral condition should have yielded more Spatial Proximal and Final errors than the Homolateral condition, because two sides are to be encoded in the former and only one in the latter condition. Both predictions were found to be false: Single and Homolateral conditions had virtually identical Side error rates (Figures 4A, 5), and Heterolateral and Homolateral conditions had virtually identical Spatial Proximal and Final-Position error rates (Figures 4A, 7). Hence such evidence strongly favours the hypothesis that side and content of the action are stored in separate STM systems, namely the spatial and the motor STM subsystems (see Baddeley, 2010; Rumiati & Tessari, 2002; Tessari & Rumiati, 2002; Ottoboni, Ceciliani, Tessari, 2021).

## Conclusion

The most interesting results of the present study concern the putative roles of “mirror” vs “anatomical” imitation, as well as of Working Memory (WM), as a function of children’s age. We believe that WM development is a good candidate to explain why performance generally improved with age (Figure 1) and why “extra-difficulties” on Side errors were strongly reduced at 9-10 years (Figure 5A). Indeed, WM includes both storage capacity and the possibility to manage between-codes conflicts, which are two of the possible causes of extra-difficulty experienced by younger children. While we did not measure WM in this study, previous literature assumes it plays a crucial role in determining the ability to imitate (Cubelli et al., 2000; Ottoboni et al., 2021; Rumiati & Tessari, 2002; Tessari & Rumiati, 2002). Age is one of the most relevant factors in influencing WM capacity since childhood, and the ability to maintain information for brief periods increases with age (the massive expansion occurs between 4 and 14 years of age; (Gathercole, 1999; Isaacs & Vargha-Khadem, 1989; Wilson et al., 1987). The steepest increase in memory performance occurs from four to eight years of age, and then becomes gradual until 11 or 12 years, where the curve flattens (Nelson, 1995). Moreover, different STM subsystems also have different developmental trajectories (Gathercole et al., 2004). Thus a (WM-grounded) increase in imitation performance with age was expected. Both GOADI and dual-routes model highlight the crucial role of cognitive resources and WM capacities. Cognitive development during childhood has been associated with the prefrontal cortex’s development, one of the last brain regions to mature (Dempster, 1992) (Casey et al., 2000; Luna et al., 2001), and this maturation is accompanied by faster information processing and an increase in reasoning ability and short-term memory capacity (Dempster, 1981; Fry & Hale, 2000; Hale, 1990).

However, WM is certainly not the only factor at play, given the surprising increase in Side error rate with age. To explain this pattern, we hypothesized a shift from purely “mirror” (or “specular”) imitation in younger (6-8 years old) children to the presence of some “anatomical” imitation in older (9-10 years old) children. Specular imitation (a type of allocentric imitation, (Bianchi et al., 2014) is more natural than anatomic (egocentric) imitation until 10 years of age (Bergès & Lézine, 1963; Brass et al., 2000; Gleissner et al., 2000; Schofield, 1976; Wapner & Cirillo, 1968). Even adults show the tendency to imitate specularly in many circumstances, as anatomical imitation is a more difficult mental operation. Indeed, anatomical imitation requires inhibiting the automatic tendency to mirror the model’s movements and an additional spatial transformation of the perceived movements from the model’s body to the imitator’s body (see patients with frontal lesions, (Chiavarino et al., 2007) or healthy adults, (Avikainen et al., 2003; Mengotti et al., 2012, 2013).

Nine- to ten-year-old children do show the prerequisites for performing anatomical imitation: they have both the required WM capacity and sufficient knowledge of the human body. As to the latter, the development of multisensory processing for own body perception seems to follow a protracted time course (Begum Ali et al., 2014; Bremner et al., 2013; Cowie et al., 2013; Nardini et al., 2013; Pagel et al., 2009)

In particular, as demonstrated using the rubber hand illusion paradigm, the visual-proprioceptive processing of own hands is strongly influenced by (the proprioceptive information of) hand position for children 4 to 9 years old (Cowie et al., 2016), but the contribution of vision relative to proprioception in hand localization is down-weighted by 10 or 11 years in comparison with younger children. Another study showed how the body schema develops progressively until the age of 8 and beliefs concerning another body mature later, from 8 to 10 years old (Assaiante et al., 2014). Moreover, eight years is the age when children develop a proper semantic knowledge of both the upper and lower body parts (Auclair & Jambaqué, 2015), but, in order to be able to use it to imitate actions in an anatomical way, higher WM capacity are needed, and this might explain why only 9-10-year-olds (the ones with higher WM capacity in our experimental group) showed such behaviour. However, provided that 9-10-year-olds *can* imitate anatomically, why did they do so, given that anatomical imitation is a more difficult cognitive operation (and was not required by the instructions)? Perhaps they could begin to “experience” and “test” their new knowledge about the body. We leave this open question, which definitely deserves further research work.

## Endnotes

1. In order to partial out the “cumulative probability” factor from the Heterolateral vs Single-effector comparison, the mathematical “removal” of the cumulative-probability factor from the Heterolateral error rate (via the inverse transformation) is not the only solution: one might mathematically “add” the cumulative-probability factor to the Single-effector error rate (via the parabolic transformation 2*x*-*x*^2^), and then compare it to the untransformed Heterolateral error rate. However, a dedicated Monte Carlo simulation study showed that the inverse transformation strategy provides slightly better estimates (with a smaller bias and therefore closer-to-nominal alpha probabilities) than the parabolic one, so we used the inverse transformation logic in all the GEE analyses.
2. We could no test the Condition x Age term directly, as this yielded a singular Hessian matrix.

## Appendix A – finding the age at which scores stabilize (or begin to rise)

Identifying the age at which Error Rate (ER) reaches a minimum or stabilization level is not a trivial task. Pairwise statistical comparisons between consecutive age groups, searching for a non-significant difference that would correspond to the stabilization point, is little reliable (as dedicated Monte Carlo simulations showed). Indeed, even if there is no flattening at all in the studied age range, but a weak, progressive decrease of ER, the probability of finding a false negative (a non-significant contrast between two consecutive age groups) is very high, and this would occur anywhere along the Age scale. Thus, we reasoned that a reliable technique should not be based on pairwise comparisons but rather on accurately estimating the whole curve linking Age to ER. Given that ERs are on a bounded 0-1 scale, effects by Age cannot generally be linear. A standard way to model such effects is to fit a sigmoidal curve. The obvious candidate would be a logistic function, with an inflexion point at ER=0.5, a lower asymptote at ER=0, and an upper asymptote at ER=1. However, ER certainly has a physiological non-zero lower limit. The upper limit cannot be safely assumed to be 1, because there might be gestures for which errors are virtually impossible, especially in some categories. Thus the 0 lower asymptote was replaced by a parameter *fl* (“floor”), and the 1 upper asymptote was replaced, for mathematical comfortableness, with *fl* + *vr*, where *vr* is the vertical range (the vertical distance between the upper and the lower asymptote). Hence, the classical logistic model, ER = 1/(1+exp(-(*s*\*Age+*i*))), with *s*=slope and *i*=intercept, became:

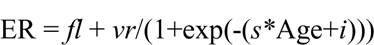

However, we are not really interested in slope ad intercept as parameters: the purpose of the present analyses is to understand at what age ER flattens after an initial drop (note that ER can also rise, and in this case, we are interested in the Age at which Error Rates begins to rise; the geometry is perfectly left-right symmetrical with respect to the ER-drop case, so we focused on Age-related drops in the following explanation). Clearly, the drop reaches the lower asymptote only at infinity, so one must set an ER level that is very close to the lower asymptote to consider the remaining part of the curve as flat. How close is “very close?” We conventionally set the threshold at 5% of the overall distance (*vr*) from the lower to the upper asymptote. Thus, the threshold ER was T = *fl*+0.05\**vr*. For instance (Fig. A1), if the lower asymptote was estimated to be *fl*=0.1, and the upper asymptote to be 0.7 (*vr*=0.7-0.1=0.6), stabilization is considered to have been reached at the Age *w* where ER equals the threshold T = 0.1+0.05*0.6=0.13.

So, we had to reparametrize the Equation so as to have *w* – the Age value where the curve intersects threshold T – as a parameter. Reparameterization led to the replacement of intercept *i* with –ln((1-0.05)/0.05)-*w*\**s*; hence the final regression equation became:

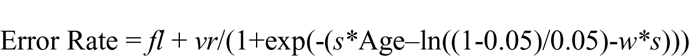

Fig. A1 shows the curve with parameters *fl*=0.1, *vr*=0.6, *s*=-2 and *w*=10.

**Figure A1.**
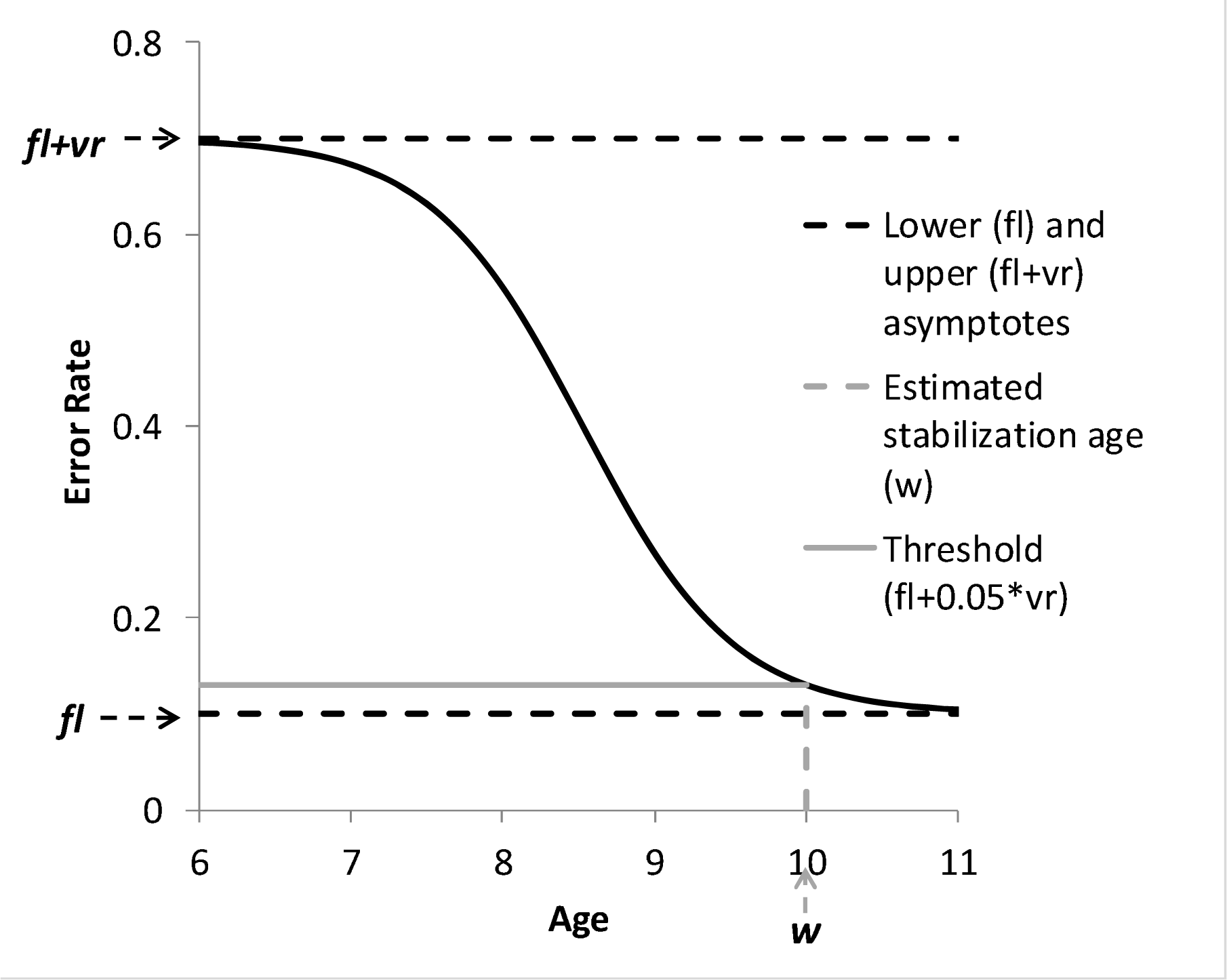
The modified logistic law (black solid curve) used to estimate the Age (w) at which error rate stabilizes. The logistic law is obtained by nonlinear regression on empirical data. Dashed black lines: upper asymptote (floor, parameter fl) and lower asymptote (ceiling, sum of parameters fl and vr). Parameter vr (vertical range) is the distance between the asymptotes. Grey solid line (threshold) is the Error Rate that is 5% of the way between the lower and the upper asymptote, and which identifies the Age were the curve (conventionally) flattens (parameter w, dashed grey segment). If the curve increases, the same geometry leads to the estimation of the conventional point where the curve begins to rise.

Sometimes this modified logistic model (which was fitted with non-linear regression in R, nls function) failed to converge. In those cases, we replaced it with a modified exponential decay model (only useful for ER *dropping* with Age). Here the general formula is ER = exp(-*s*\*(Age)). Since the top score (ER=1) can only be reached at Age=0, we replaced Age with Age-6 (the youngest participants were 6-year-old children), so that the equation could fit the full range of ERs. Again we implemented a lower physiological limit that can be higher than zero, *fl*, and a vertical range *vr*, as free parameters:

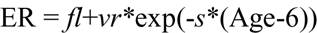

Reparametrization, with Age-6 value *w* at which Error Rate reaches the threshold T, led to *s* = −ln(0.05)/*w*; hence the final equation:

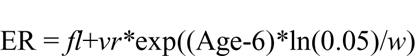

An example is shown in Figure A2, showing the curve with parameters *fl*=0.1, *vr*=0.6, and *w*=4 (corresponding to Age=10).

**Figure A2.**
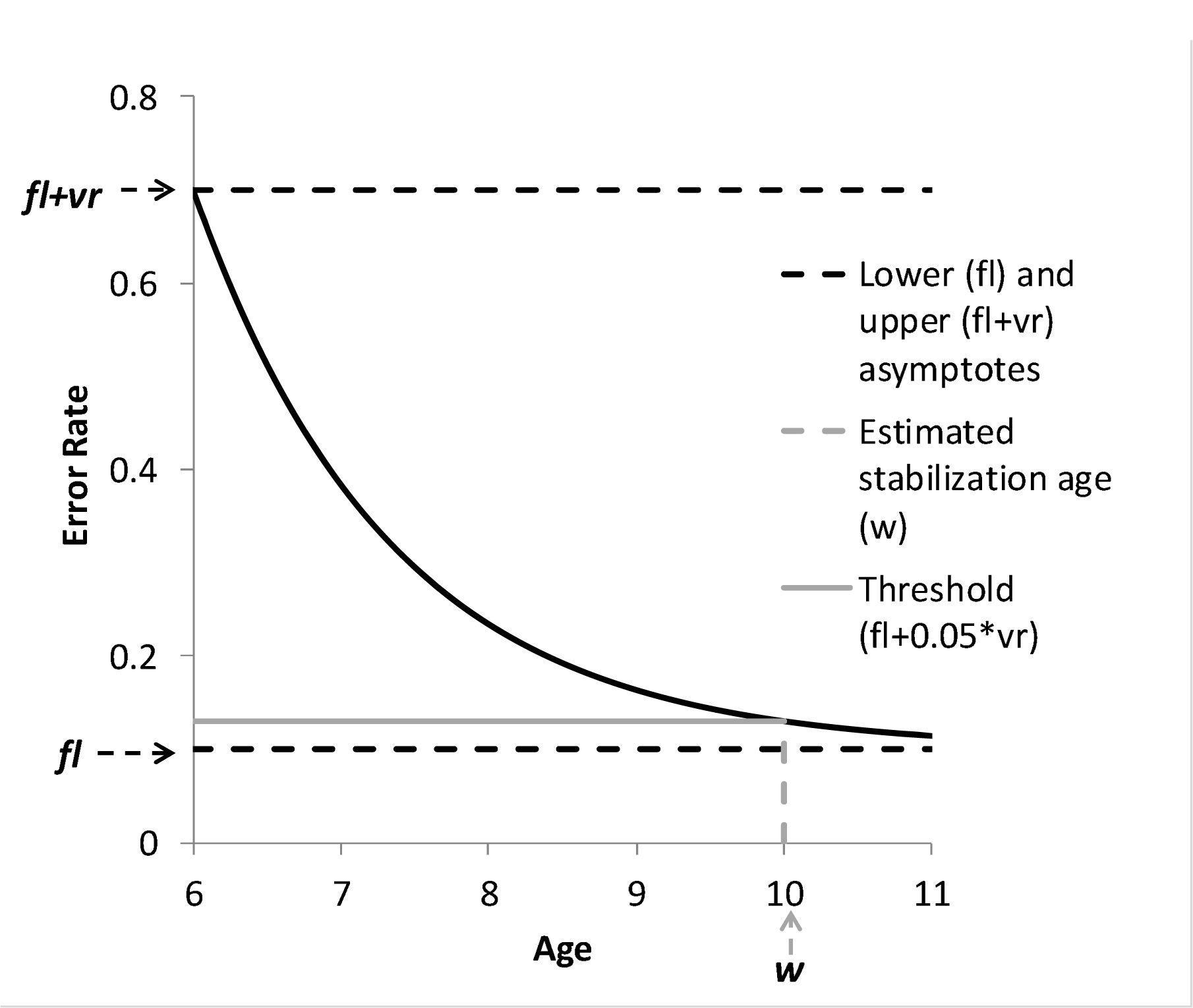
The modified exponential decay law. See Fig. A1 for conventions.

## Appendix B – the inverse transformation

### Area of application of the inverse transformation

Formally, the inverse transformation *x* = 1-(1-*y*)^1/2^ holds under the statistical assumption that the probabilities of mis-processing the arm and of mis-processing the leg are identical; empirically, the transformation holds even though the two probabilities show huge differences: Monte-Carlo simulation work showed that with a .26 difference in baseline error probability for arm and leg (.05 vs .31), the adjustment by the inverse transformation is off by about .01. Anyhow, we could reject the hypothesis that there were sizeable differences by comparing the Single-effector trials employing an arm vs a leg by GMLM for the three error categories that underwent the inverse transformation. No significant differences were found: Side errors, Wald χ^2^=2.272, p=.132 (Arm=.086, Leg=.108); Spatial Proximal errors, Wald χ^2^=.033, p=.856 (Arm=.045, Leg=.039); Spatial Final errors, Wald χ^2^=.373, p=.542 (Arm=.013, Leg=.001).

### The theoretical value of the inverse transformation

By its very nature, the inverse transformation performs a perfect cancellation of the disadvantage of the Heterolateral with respect to the Single-effector condition, when in the former condition, the probability of an error on the arm is statistically independent of the probability of an error on the leg. Interestingly, STM/WM limits, conflict managing limits, and anatomical imitation strategy (if applied to the whole body on each trial) all produce a violation of that statistical independence, thus becoming “visible” as under-corrections or over-corrections by the inverse transformation – hence the value of this mathematical technique. These premises were all confirmed by dedicated Monte Carlo simulations (N=10,000 for each sample), and are detailed below.

STM/WM limits produce violations of independence between arm and leg error probabilities by boosting the relative frequency of side errors made on a single limb: by WM/STM limits alone, subjects are likely to forget the side of a single limb, but much less likely to forget the sides of both limbs. Overall, this produces a surplus of errors that the inverse transformation under-corrects, so the adjusted Heterolateral Side error rate would turn out to be higher than the Single-effector Side error rate. We referred to this as “extra-difficulties” in the main text.

A virtually identical pattern is predicted by putative “conflict-managing” limits, according to which the side of one limb is mistaken as the side of the other limb (e.g., left arm and right leg becomes left arm and left leg) because the child is not able to manage the conflict between the spatial labels (“left” vs “right”). The extra-errors due to this phenomenon would all (or mostly) regard a single limb, and the result would again be an under-correction by the inverse transformation.

The exception to this rule is the hypothesis of an anatomical imitation strategy in which the participant reverses the whole body’s side codes. Here, side errors would either regard both limbs simultaneously, or neither of them. Overall, the inverse transformation would over-correct, and the adjusted Heterolateral scores would turn out to be *smaller* than the Single-effector score.

Importantly, the hypothesis that the anatomical imitation strategy is not applied to the whole body, but rather, separately and in a statistically independent way, to the arm or the leg, predicts neither over-nor under-correction: the Heterolateral error rate increase would be perfectly cancelled by the inverse transformation, and the adjusted Heterolateral score would match the Single-effector score.

In older (9-10) children, who made many more Side errors than younger ones (a result which we interpreted in terms of an anatomical imitation strategy), the adjusted Heterolateral and Single-effector scores were indeed very similar (Fig. 5A). The most parsimonious account – albeit counterintuitive-is that they had neither STM/WM limitation nor interference phenomena and applied an anatomical imitation strategy separately on arm and leg, which predicts the empirically observed pattern. However, this conclusion is premature, not only because it is based on a non-significant result, but because it might have been caused by two opposite effects that cancelled each other out: if children’s performance had been affected by both (e.g.) conflict-managing limits causing an under-correction, and a whole-body anatomical imitation strategy causing an over-correction, the combined result would have been that of an almost perfect correction – matched Single-effector and adjusted-Heterolateral scores, which was in fact observed. To the best of our knowledge, there is no way to tell which interpretation is correct, at least, not on the grounds of the present experiment: future research will address this interesting issue.

The possibility that older children might have applied an anatomical imitation strategy to the arm and not to the leg (or vice versa) within the same trial is interesting in its own right. Clearly, mental rotation would clash with this idea; however, anatomical imitation might not imply internal analogical rotations, but rather, only changes in abstract, spatial labels, possibly separate for arm and leg.

## Footnotes

1 The real number of triple errors was 6; however, in the Table each of those errors was counted twice (in the negative direction: −12 overall), because, when subtracting triple errors from the overall set of trials, one has to consider that there are _two_ “extra” errors in each triple that need to be subtracted in order to obtain a single error (a single trial) and make the computations come out right.

